# Caves as species pumps: key innovations, isolation, and periodic introgression drive the world’s largest cavefish radiation in a dynamic karstic landscape

**DOI:** 10.1101/2024.09.12.612638

**Authors:** Tingru Mao, Yewei Liu, Mariana M. Vasconcellos, Shipeng Zhou, Gajaba Ellepola, Jian Yang, Marcio R. Pie, Madhava Meegaskumbura

## Abstract

Species diversification is shaped by intricate interactions among biotic drivers, including gene flow, hybridization, and key innovations, and abiotic drivers, such as historical climate change, geological events, and ecological opportunity. However, the relative contributions of these drivers in large radiations remain poorly understood. We investigate the interplay among these factors in the diversification of *Sinocyclocheilus*, a cavefish radiation comprising 79 species. *Sinocyclocheilus* include typical surface-dwelling forms, with well-developed eyes and pigmentation, to cave-dwelling forms with regressed eyes, reduced pigmentation, and the presence of a horn and a hump. Using reduced representation genomic data (RADseq), we show extensive gene flow events across different species, with introgression playing a key role compared to incomplete lineage sorting in creating phylogenetic discordance and contributing genetic variation for cave adaptation and diversification in this group. Key traits such as eye degeneration, reduced pigmentation, and horn evolved independently multiple times, as adaptations for effectively exploiting cave environments. Furthermore, the uplift of the Tibetan plateau and the late Miocene cooling also significantly impacted speciation rates. Demographic analyses suggest population expansions during the Gonghe Movement and stability during the Last Glacial Maximum, possibly due to cave refugia. Periodic events of introgression promoted by isolation and reconnections due to the changing climate and geological activity, combined with the repeated evolution of key cave-adapted traits, are the primary drivers of this radiation. Our findings underscore the complex interplay of biotic and abiotic factors in the evolution of *Sinocyclocheilus* fish, offering new insights into the mechanisms of cave adaptation and diversification.

## 1 INTRODUCTION

Lineage diversification often involves a complex interplay between biotic and abiotic drivers. For instance, the rapid diversification that results in evolutionary radiations is often attributed to a combination of biotic factors, including hybridization, introgression, and key innovations (Abbott et al., 2013; Dowling & Secor, 1997; Heard & Hauser, 1995; Hunter, 1998; Koblmüller et al., 2010; Meier et al., 2017; Seehausen, 2004; Wainwright et al., 2015) as well as abiotic factors, such as ecological opportunities (Salzburger, 2009; Stroud & Losos, 2016; Wagner et al., 2012; Yoder et al., 2010), geological events and climate change (Erwin, 2009; Favre et al., 2015; Hoorn et al., 2013; Kozak & Wiens, 2010; Linder et al., 2014; Near et al., 2012). Therefore, assessing their respective contributions remains a central goal in evolutionary studies.

As a biotic interaction, hybridization—one of the most common biotic drivers—is believed to facilitate ecological divergence and speciation by generating novel trait combinations that enable the exploitation of previously inaccessible resources (Marques et al., 2019; Seehausen, 2004). The emergence of such traits can enhance the fitness of hybrids in specific environments and enable them to colonize previously unoccupied niches (R. L. Moran et al., 2022). As a result, hybridization not only triggers adaptive radiations but also sustains them beyond the initial speciation events (Meier et al., 2017; Porretta & Canestrelli, 2023; Seehausen, 2004). Increasing evidence suggests that both ancient and contemporary introgression are widespread among diverse animal groups (Kim et al., 2022; Lavretsky et al., 2021; Suvorov et al., 2022; Taylor & Larson, 2019; Hirase et al., 2021). Some studies propose that introgression serves as a critical source of adaptive genetic variation, potentially exacerbating disparities between populations rather than impeding adaptation and diversification (Abbott et al., 2013; Ellstrand & Rieseberg, 2016; Ford et al., 2015; Wang et al., 2023). Given the prevalence of hybridization in comparison to the relative rarity of new mutations, introgression may play a significant role in shaping the evolution of populations and, ultimately, species (Abbott et al., 2013). In addition, hybridization and introgression, being dynamic and context-dependent processes, contribute not only to the formation of biodiversity patterns but also to their ongoing reshaping in response to environmental changes (Brauer et al., 2023; Ryan et al., 2018).

Key innovations also represent another important biotic driver of diversification, operating through three primary ecological mechanisms (Heard & Hauser, 1995; Ronco & Salzburger, 2021). First, a novel trait may represent an evolutionary breakthrough in function, enabling its carriers to exploit previously unfilled or unoccupied niche spaces (Near et al., 2012). Second, key innovations can significantly enhance the fitness of the organism, providing a competitive advantage by optimizing resource use or improving the evasion of predators or parasites (Farrell et al., 1991). The third ecological mechanism by which key innovations can promote diversity involves increased specialization. Such innovations expand the morphospace by facilitating new combinations of functional traits alongside existing ones, thereby directly increasing ecomorphological disparity and enabling greater species coexistence (Heard & Hauser, 1995; Ronco & Salzburger, 2021; Wainwright, 2007). The evolution of key phenotypic innovations can also create new ecological opportunity (Daane et al., 2019; Dumont et al., 2012; Hunter, 1998; Maia et al., 2013; Miller et al., 2023; Wellborn & Langerhans, 2015; Yoder et al., 2010), by allowing the exploitation of new habitats and niches or conferring a competitive edge. Therefore, key innovations and ecological opportunity are closely intertwined in driving species diversification.

Among the abiotic drivers, ecological opportunity often arises through changes in the organism ecology, resulting in reduced competition or an increase in resource availability (Schluter, 2000), and has been widely recognized as a primary driver of adaptive radiations (Losos, 2010; Rabosky, 2009; Yoder et al., 2010). Moreover, ecological opportunities may interact synergistically with other mechanisms, such as hybridization (Meier et al., 2019), potentially enhancing its impact on adaptive radiations. Understanding the interplay between these factors offers valuable insights. These rapid radiations are frequently observed in nature and have been extensively documented in phylogenetic studies across a variety of taxa, including African cichlid fishes (Seehausen, 2006), iguanian lizards (Townsend et al., 2011), passerine birds (Cai et al., 2020), and Australo-Papuan rodents (Roycroft et al., 2020). Likewise, historical climatic and geological events have been acknowledged as significant catalysts of evolutionary dynamics, promoting speciation and accelerating lineage diversification (Buckley et al., 2021; Leprieur et al., 2021; Near et al., 2012; Thomaz et al., 2015). Examples of such events include the Late Miocene cooling (11.6–5.3 Ma) (Near et al., 2012) and the Tibetan uplift (Qian et al., 2023; X. Wang et al., 2016), which had a great impact on fish biodiversity. Similarly, the Pleistocene “species pump” effect illustrates how lineages across several taxa can quickly diversify in response to climate oscillations (April et al., 2013; Schoville et al., 2012; Sholihah et al., 2021).

With the growing availability of genome-wide data for non-model organisms, it is now possible to explore the factors that drive diversification of large and, yet poorly-studied systems. The *Sinocyclocheilus* cavefishes, representing the largest cavefish radiation in the world with 79 currently described species (Shao et al., 2024), provide a unique opportunity to investigate the mechanisms underlying rapid evolutionary diversification in south and southeast Asia. While previous studies have primarily relied on mitochondrial DNA to infer phylogenetic relationships (Jiang et al., 2023; T.-R. Mao et al., 2021; Wen et al., 2022; Xiao et al., 2005), this approach can lead to incorrect or misleading conclusions (Ballard & Whitlock, 2004; Rubinoff & Holland, 2005; Toews & Brelsford, 2012), especially when introgression is present (T. Mao et al., 2022; Jacobs & Therkildsen, 2019). Considering that introgression is being increasingly recognized as a critical source of genetic variation for adaptation and diversification, it is crucial to examine its role in the diversification of *Sinocyclocheilus*. Furthermore, this genus exhibits a remarkable diversity of morphological forms, ranging from typical surface-dwelling fish with normal eyes, pigmentation, and absence of horn and hump, to cave-dwelling forms characterized by eye degeneration, reduced pigmentation, and the presence of a horn and a hump (Borowsky, 2018; Jiang et al., 2023). These cave-associated traits are likely key innovations that have promoted diversification within *Sinocyclocheilus,* contributing to their evolutionary success.

The remarkable species diversity in *Sinocyclocheilus* is also likely driven by significant geological events, particularly the dramatic uplift of the Tibetan Plateau and subsequent climatic changes, including the intensification of the Asian monsoon and increased precipitation (Z. Li et al., 2008; Ma et al., 2019; Xiao et al., 2005). The geological events, combined with the effects of the monsoonal climate, are the primary forces behind karst cave formation in southwest China (Wen et al., 2022; L. Zhang, 2012). The unique ecological conditions within these cave environments were likely a promoter of the extensive adaptive radiation in *Sinocyclocheilus*. Considering both the temporal and spatial aspects of their evolutionary history and geographic distribution, this genus represents an ideal model to investigate and quantify the impacts of the Tibetan Plateau uplift, global cooling, and intensified monsoons on its rapid evolutionary radiation. In addition, previous research has shown that the demographic history of some *Sinocyclocheilus* species, specifically *S. grahami*, *S. rhinocerous*, and *S. anshuiensis*, are also linked to paleoenvironmental events (J. Yang et al., 2016). However, the demographic history of other *Sinocyclocheilus* species and whether those exhibit a congruent response to the same paleoenvironmental events remain to be determined.

A prior study inferring the phylogenetic relationships in *Sinocyclocheilus* using genome-wide sampling (RADseq) (T. Mao et al., 2022) across 21 species has already revealed unaccounted species diversity in this group, with preliminary evidence of rapid speciation and introgression. Here, we build upon this work, increasing taxon sampling of genome-wide single nucleotide polymorphism (SNP) to 50 *Sinocyclocheilus* species to address specific hypotheses about the biotic and abiotic drivers of diversification in this group. We aim to address the relative roles of introgression, key innovations, geological events, and climate change. The specific objectives of our study are: (1) to infer a comprehensive phylogeny for the genus using RADseq data, examining the roles of introgression and incomplete lineage sorting (ILS) during their rapid diversification based on the phylogenetic discordance to the mitochondrial DNA-based phylogeny; (2) to investigate potential hybridization and introgression events among *Sinocyclocheilus* species, particularly sympatric ones, assessing their contribution to diversification; (3) to evaluate the impact of the uplift of the Tibetan Plateau, global cooling, and the intensity of the East Asian monsoon on the speciation rates of *Sinocyclocheilus* by applying macroevolutionary models of diversification dynamics; (4) to determine whether four cave-associated traits — eye degeneration, horn, hump, and reduced pigmentation—are key innovations linked to increased diversification, using macroevolutionary models to reconstruct their ancestral states and trace their evolutionary history; and (5) to reconstruct the demographic history of well-sampled species, examining how they have responded to paleogeographic and climatic changes over time.

## 2 MATERIALS AND METHODS

### 2.1 Taxon sampling

To investigate the abiotic and biotic factors driving the diversification of *Sinocyclocheilus*, we conducted a comprehensive phylogenomic analysis of the group. For this, between 2017 and 2020, we undertook extensive field collections of *Sinocyclocheilus* fish across the Guangxi, Yunnan, and Guizhou provinces in China (Figure S1). Due to the rarity and challenges of acquiring *Sinocyclocheilus* samples, live specimens were collected and fin clips were obtained without destructive sampling. The fin clips were immediately frozen at −85 °C for DNA preservation. We sampled 257 specimens across 50 species, including 48 recognized *Sinocyclocheilus* species and two potential cryptic species (*S*. cf. *cyphotergous and S.* cf*. guanyangensis*). We also included six specimens from two outgroup species: *Cyprinus carpio* and *Puntius semifasciolatus* (Table S1).

### 2.2 RAD library construction and sequencing

Details concerning DNA extraction, library preparation, sequencing protocols, and raw data filtering procedures are provided in the Supplementary Material.

### 2.3 RADseq data assembly

We used ipyrad version 0.9.56 (Eaton & Overcast, 2020) to process the demultiplexed reads and assemble them using a *de novo* method with the same parameter settings as in T. Mao et al. (2022). Detailed parameter settings are provided in the Supplementary Material. We generated multiple datasets for different analyses (Table S2), and a flowchart summarizing the key analytical steps and bioinformatics methods used in this study is provided in Figure S2.

The suitability of the clustering threshold was assessed heuristically by executing steps 3 and 4 of the ipyrad pipeline and examining the hetero_est and error_est metrics. This evaluation ensured that the error rate (approximately 0.1%) and heterozygosity estimates (ranging from 0.001 to 0.05) remained within acceptable limits to prevent erroneous over-clustering or over-splitting of loci. To mitigate the potential confounding effects of paralogs, we employed the R package *Phrynomics* v2.0 (Leache et al., 2015) to remove any non-binary single nucleotide polymorphisms (SNPs).

### 2.4 Phylogenetic inference

To investigate phylogenetic discordance, we employed multiple phylogenetic inference methods. These included a maximum likelihood approach with IQ-TREE2 (Minh et al., 2020), the multispecies coalescent-based method SVDquartets (Chifman & Kubatko, 2014), and a gene tree summarization approach with ASTRAL-III (C. Zhang et al., 2018). The ASTRAL score, representing the proportion of induced quartet trees from the input set present in the species tree, serves as a measure to assess the extent of incomplete lineage sorting (ILS). To additionally visualize the overall phylogenetic structure of our SNP data, including reticulate relationships, we constructed a phylogenetic network using the complete matrix of all individuals with the NeighborNet algorithm in SplitsTree 4 (Huson & Bryant, 2006). To infer the phylogenetic discrepancies between nuclear DNA (nDNA) and mitochondrial DNA (mtDNA), CYTB and ND4 sequence fragments were obtained for 63 *Sinocyclocheilus* species and two outgroup species from GenBank (Table S3) to construct a mitochondrial DNA tree. *Cyprinus carpio* and *Puntius semifasciolatus* were included as outgroup species. Further details about the phylogenetic inference methods are given in Supplementary Material.

### 2.5 Tests of introgression and detection of gene flow

To assess the impact of hybridization and ILS on topological discordance, we conducted MSCquartets analyses (Rhodes et al., 2021). To further investigate introgression among species, we applied Patterson’s D statistic (ABBA-BABA test), along with the f4-ratio statistic (Patterson et al., 2012) and its f-branch metric (Malinsky et al., 2018), using the Dsuite v0.5 software package (Malinsky et al., 2021). The direction and relative magnitude of gene flow among species and lineages were estimated with TreeMix v.1.13 (Pickrell & Pritchard, 2012) as implemented in ipyrad. Furthermore, to better clarify the reticulate evolutionary relationships within *Sinocyclocheilus*, we constructed species networks using SNaQ (Solís-Lemus & Ané, 2016) implemented in the Julia package PhyloNetworks v0.15.3 (Solís-Lemus et al., 2017), which accounts for both ILS and gene flow in estimating a phylogenetic network.

Specimens of *S. cyphotergous* and *S. punctatus*, both belonging to clade D, were collected from the same cave. Notably, the individuals Liu129 and Liu354 exhibited morphological traits distinct from those of *S. cyphotergous* and *S. punctatus*, displaying intermediate phenotypes, which suggests they may be hybrids of these two species. To elucidate the genetic composition and heterozygosity levels of Liu129 and Liu354, we employed the Bayesian clustering software STRUCTURE v.2.3.4 (Pritchard et al., 2000), without presupposing the identities of the *S. cyphotergous* and *S. punctatus*. Further details on the analyses of introgression and hybridization are provided in the Supplementary Material.

### 2.6 Divergence time estimation and dynamics of diversification

The time-calibrated phylogenetic tree was reconstructed using SNAPP v.1.4.1 (Bryant et al., 2012), as implemented in BEAST v.2.7.3 (Bouckaert et al., 2014), following the methodology described in Stange et al. (2018). Due to the absence of reliable fossil records for the *Sinocyclocheilus* genus, tree calibration was based on four secondary calibration points derived from R. Li et al. (2021) (Supplementary Material for details). The resulting maximum clade credibility (MCC) tree was pruned to include the 48 currently recognized *Sinocyclocheilus* species for subsequent analyses. Additional details about the estimation of divergence times are provided in the Supplementary Material.

To investigate the temporal dynamics of lineage accumulation in *Sinocyclocheilus*, we first produced a lineage-through-time (LTT) plot using the R package phytools 1.2-0 (Revell, 2012). Then, we employed a series of macroevolutionary models to elucidate the relative contribution of temporally varying abiotic factors driving the diversification dynamics of *Sinocyclocheilus*. These models were fitted to our calibrated phylogeny using a Maximum Likelihood framework (Condamine et al., 2013; Morlon et al., 2011, 2016) in the R package RPANDA (Morlon et al., 2016). We developed seven paleoenvironment-dependent models, including the impact of global paleotemperatures, East Asian monsoons, and paleoelevation of the Tibetan Plateau on speciation rates, and nine time-dependent diversification models to explore the association between diversification rate shifts and environmental changes. In addition, we tested the effect of diversity dependence on the diversification history of *Sinocyclocheilus* using the R package DDD v4.2 (Etienne & Haegeman, 2012). The details of the diversification dynamics analyses are described in the Supplementary Material. All models were evaluated and ranked based on the corrected Akaike Information Criterion (AICc) and Akaike weights (AICw).

To determine whether cave-associated traits in *Sinocyclocheilus* are linked to an increase in diversification rates, we investigated state-dependent diversification, focusing on four putative key innovations: degenerated eye, horn, hump, and reduced pigmentation. We used the Hidden State Speciation and Extinction (HiSSE) framework in the corresponding R package v4.3.2 (Beaulieu & O’Meara, 2016) evaluating 30 models following the comprehensive model-testing approach of Blaimer et al. (2023). Model specifications and parameters are detailed in the Supplementary Material. To accurately reflect the occurrence of these traits within *Sinocyclocheilus*, we calculated sampling fractions for character states, specifying the proportion of sampled versus unsampled trait occurrences (Table S4). Model performance was again assessed using the AICc and AICw, with diversification rate estimates and trait reconstructions visually summarized for the top-performing models.

### 2.7 Ancestral state reconstruction of traits and demographic history inferences

Stochastic character mapping (Bollback, 2006) was performed using the R package phytools (Revell, 2012) to investigate the evolution of four cave-associated traits: degenerated eye, horn, hump, and reduced pigmentation in *Sinocyclocheilus*. Details about ancestral state reconstruction analyses are provided in the Supplementary Material.

In addition, we reconstructed the demographic history for 15 *Sinocyclocheilus* species with at least seven individuals using Stairway Plot v.2.1.1 (X. Liu & Fu, 2020). This software processes folded SNP frequency spectra not requiring ancestral alleles or a reference genome, making it particularly suitable for non-model organisms. The specific descriptions of the demographic inference analyses are detailed in the Supplementary Material.

## 3 RESULTS

### 3.1 Phylogenetic inference

We obtained, on average, ∼3.3 billion raw reads per sample, with an average error rate of 0.03%. Summary statistics and additional information on the raw sequencing results are provided in Table S5.

The concatenation method, as implemented in IQ-TREE (Figure 1A), along with two coalescent-based strategies — SVDquartets (Figure S3A) and ASTRAL (Figure S3B) — clearly delineated six distinct clades (A, B, C, D, E, and F), establishing a well-supported phylogenetic backbone for *Sinocyclocheilus*. With a concatenated dataset of 9,057 loci, IQ-TREE produced a well-resolved phylogeny, although three nodes had bootstrap support values under 80% (Figure 1A). In comparison, the topology generated by SVDquartets revealed ten nodes with bootstrap support values below 80% (Figure S3A), while the species tree inferred by ASTRAL displayed exceptionally high support for all nodes (local posterior probability = 1) (Figure S3B). Coalescent-based methods like SVDquartets and ASTRAL detected several topological discrepancies when compared to the trees using a concatenation approach: two inconsistencies were observed with SVDquartets, while six were identified with ASTRAL (Figure S3A, Figure S3B). The quartet-based species tree estimated in ASTRAL was derived from 1,010,733,531,834 induced quartet gene trees, representing 64.02% of all quartets in the species tree. The normalized quartet score of 0.64 indicates substantial discordance among gene trees, suggesting the presence of incomplete lineage sorting (ILS).

**FIGURE 1.**
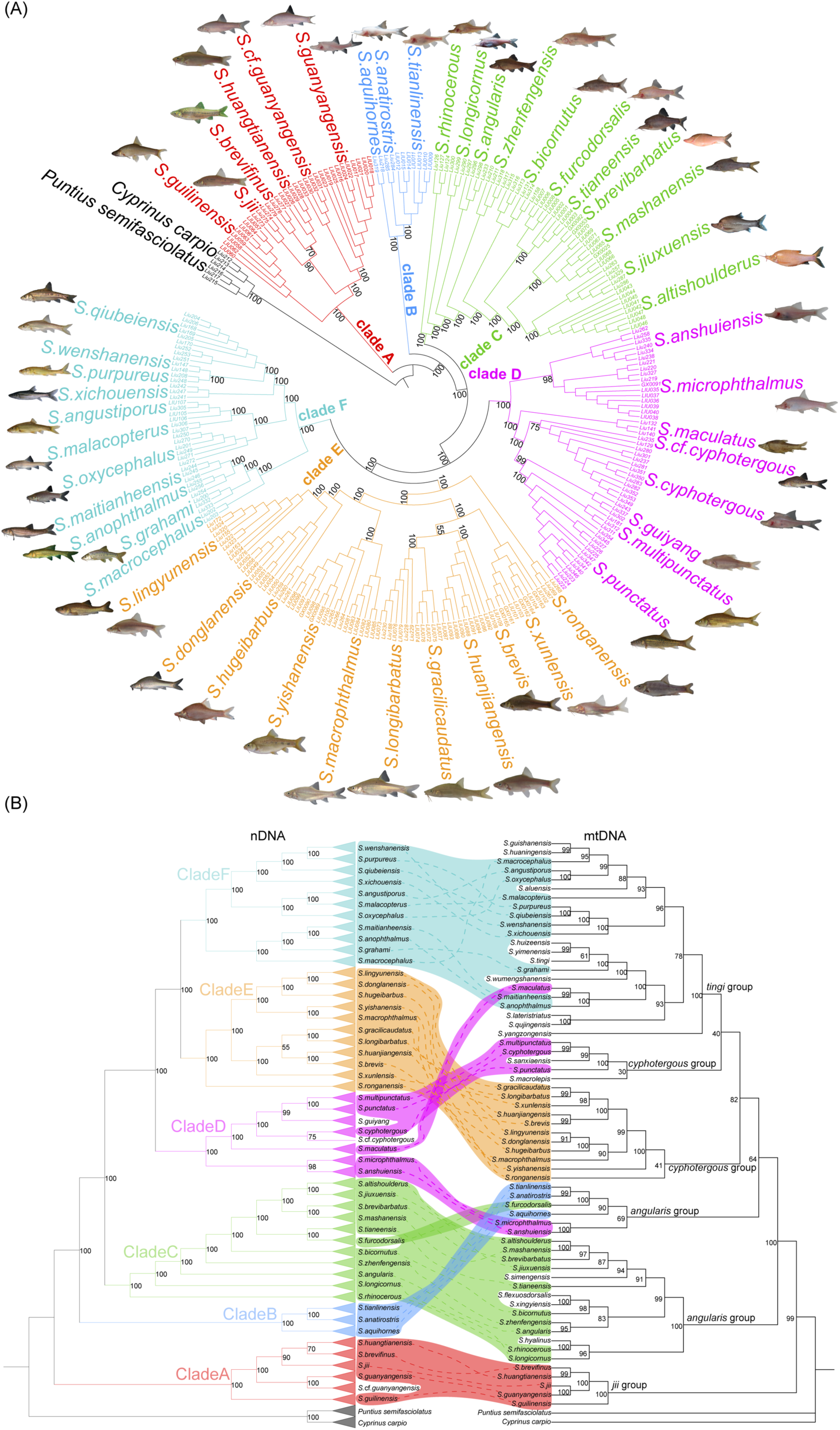
Phylogenetic trees of *Sinocyclocheilus* cavefish based on RADseq and mtDNA data inferred by IQ-TREE with *Cyprinus carpio* and *Puntius semifasciolatus* as outgroups. (A) Phylogeny based on 9,057 concatenated RAD loci. Node numbers indicate ultrafast bootstrap support values. (B) Incongruencies between the nuclear RADseq loci (left) and the mitochondrial DNA tree (CYTB and ND4) (right). Ultrafast bootstrap values are displayed on branches of both trees. Connectors highlight phylogenetic discordances. The four taxonomic classifications (jii group, angularis group, cyphotergous group, and tingi group) are based on morphological characteristics as established by Zhao & Zhang, (2009).

The phylogenetic tree derived from mtDNA data revealed weak support in the backbone structure and showed substantial topological inconsistencies compared to nDNA trees (Figure 1B). The observed mito-nuclear discordance underscores the complex relationships among clades and the intricate connections among species within these clades, challenging the notion that a simple, linear branching process can fully encapsulate their evolutionary history. Our Neighbor-Net results further corroborate these findings, displaying widespread reticulation events within the genus, further indicating significant topological conflicts (Figure S3C). Additionally, the analysis of quartet concordance factors (qcCFs) underscores the role of hybridization as a substantial factor contributing to the discordance observed in mito-nuclear gene trees (Figure S4). Specifically, about 15.38% of tests employing the T3 model rejected the multispecies coalescent (MSC) hypothesis at the α = 0.01 significance level. Simplex plots (Figure S4) show that the blue circles are closer to the vertex across the *Sinocyclocheilus* phylogeny, indicating that incomplete lineage sorting (ILS) played a relatively minor role. Instead, other mechanisms such as introgression were more likely within *Sinocyclocheilus*.

### 3.2 Complex patterns of hybridization and introgression within Sinocyclocheilus

We identified a significant signal of introgression among most species pairs using the ABBA – BABA statistic in Dsuite, with 4,168 out of 19,600 trios exhibiting a significant D value at P < 0.05 following standard block-jackknife procedures (Table S6). These findings provide compelling evidence of widespread historical introgression across *Sinocyclocheilus* evolutionary history. Based on D-statistics, we detected the strongest signal of excess allele sharing between the *S. lingyunensis* and *S. microphthalmus* (D = 0.57, Z-score = 27.46, P < 0.001; Table S6, Figure 2A). Remarkably, despite belonging to different clades (clades E and D, respectively), these two species occur in the same underground river. Additionally, *S. guilinensis* showed excess allele sharing with all species except those from clade A. There was also an excess of allele sharing between clades E and F (Figure 2A), indicative of ancestral hybridization. High gene flow between several species pairs was supported by the f4-ratio test (Z > 3, P < 0.001, f4-ratio > 0.1; Table 1; Figure 2B). The f-branch test also indicated multiple instances of introgression (Figure 2C), with *S. macrocephalus* and *S. oxycephalus* exhibiting the most compelling evidence (fb = 0.26), likely due to their cohabitation in the same cave and the occurrence of hybridization between them. Most instances of gene flow appear to be recent and occur between closely related species within clades, especially in clade E (Figure 2). In summary, Dsuite analyses revealed widespread introgression, suggesting multiple hybridization events among ancestral lineages, as well as among contemporary species (Figure 2, Table S6).

**FIGURE 2.**
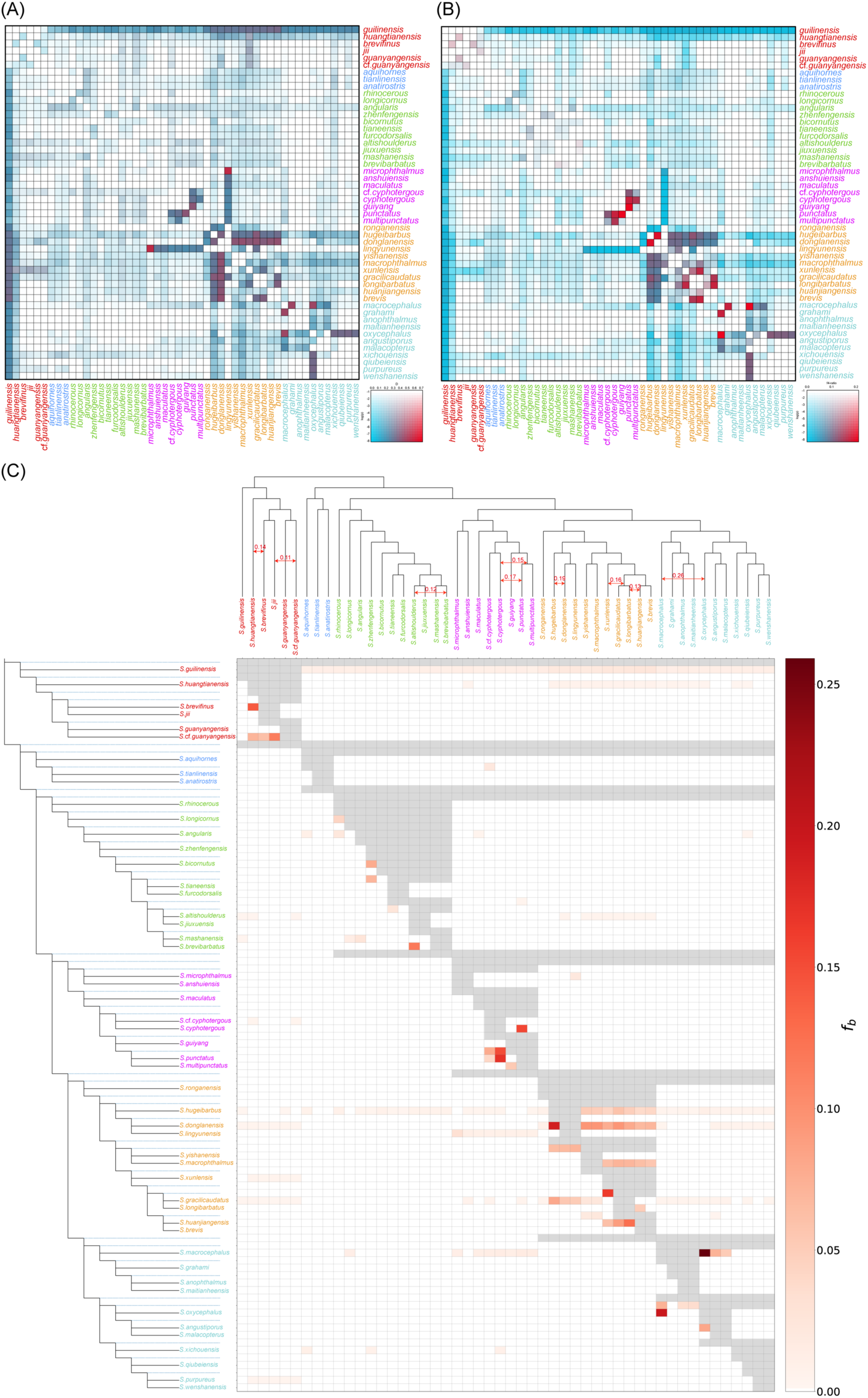
D-suite statistics results. (A) Heatmap of D-statistics estimates indicating significant introgression across most species. (B) Heatmap of f4-statistical values illustrating the proportion of the genome influenced by admixture. (C) Heatmap of f-branch statistic (fb) values tested via Dsuite using the topology recovered by SVDquartets. Detection of gene flow via the f-branch method is depicted, where dotted lines in the phylogeny indicate ancestral lineages and solid lines represent extant lineages. The f-branch statistics reveal extensive introgression across taxa and lineages. Grey cells denote combinations where f-branch values could not be calculated due to the constraints of the tree topology. High values above 0.1 are indicated in the matrix and visualized as red double-ended arrows on the top dendrogram.

**TABLE 1.**
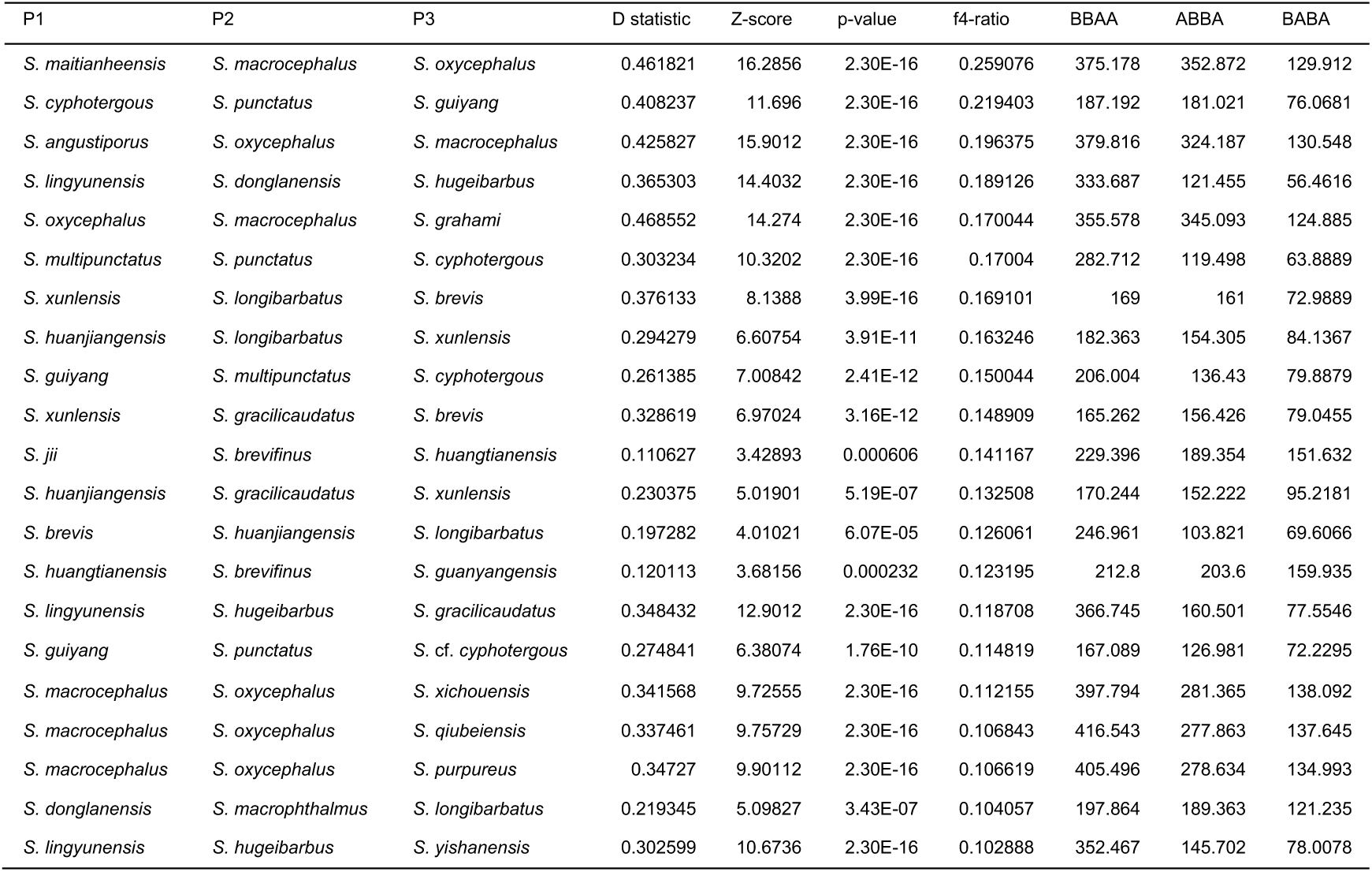
Significant results of the Patterson’s D statistic (ABBA-BABA test) and the f4-ratio test (Z scores > 3, P < 0.001, f4-ratio > 0.1).

Our Treemix analysis also detected many gene flow events within and between clades (Figure S5), with several instances of ancestral gene flow between different clades (Figure S5A). Similarly, the PhyloNetworks analysis identified substantial gene flow within clades, particularly ancestral gene flow occurring in clade A, C, and E (Figure S6). Although some of the introgression events detected differed between PhyloNetworks, TreeMix, and the D-statistic, several events were consistently detected across all three methods. These include the introgression between *S. altishoulderus* and *S. brevibarbatus* in clade C, between *S. punctatus* and *S. cyphotergous* in clade D, and between *S. oxycephalus* and *S. macrocephalus* in Clade F (Figure 2, Figure S5, Figure S6). Additionally, the Structure analysis provides evidence of admixture between *S. punctatus* (P2) and *S. cyphotergous* (P3). Individuals Liu129 and Liu354, which have intermediate phenotypes, were supported as genetic intermediates (Figure 3).

**FIGURE 3.**
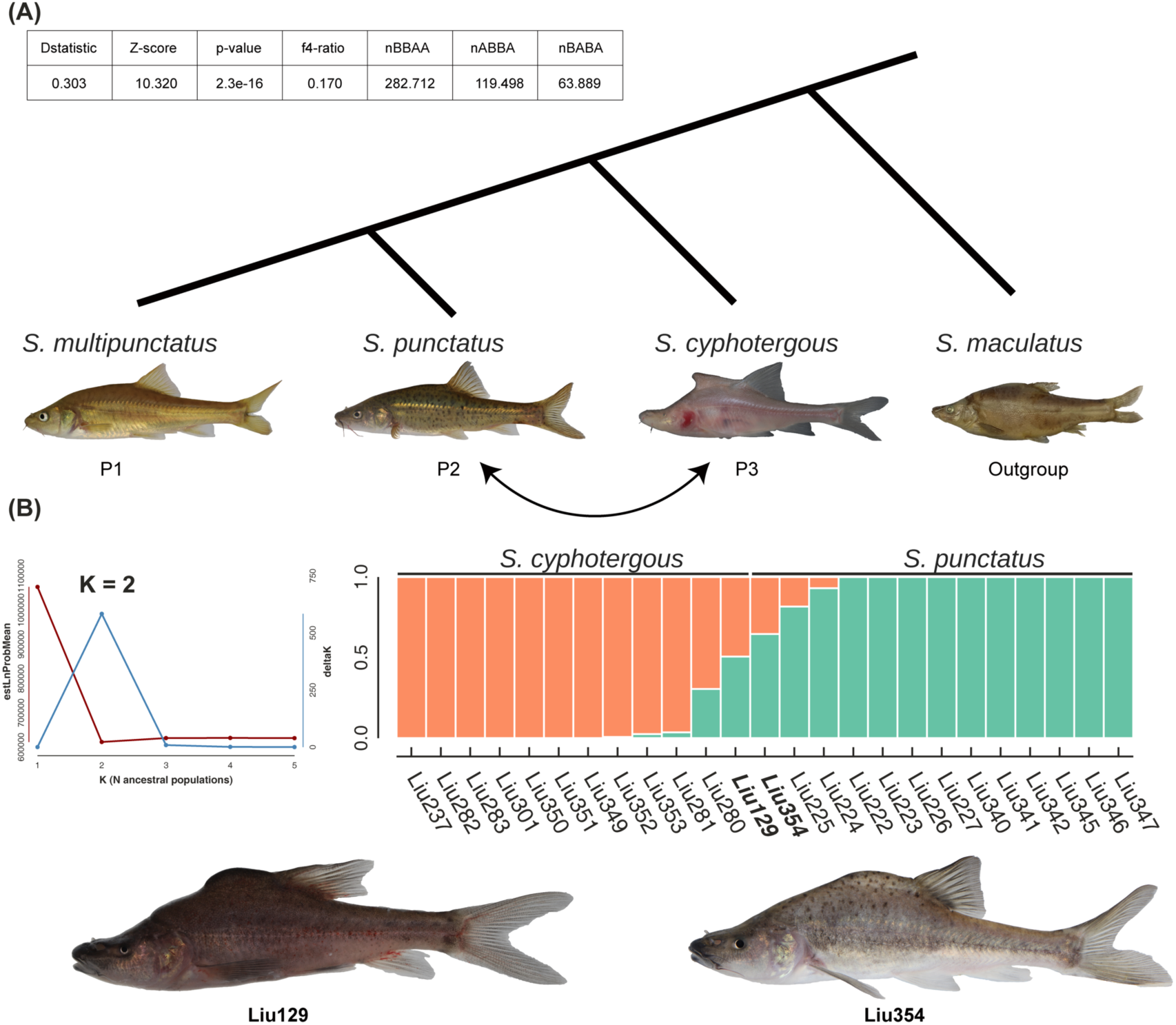
Results of ABBA-BABA test and Structure analysis. (A) ABBA/BABA result. P1, P2, P3 and outgroup were represented in order. Curved double-headed arrow indicates introgression event. (B) Results of Structure analysis for *S. cyphotergous* and *S. punctatus* complex groups. The number of populations inferred by plotting the mean log probability (red) and delta K (blue) suggests that two populations is the most likely result. Samples Liu129 and Liu354 are hybrid individuals of the species *S. cyphotergous* and *S. punctatus*, respectively.

### 3.3 Diversification dynamics in *Sinocyclocheilus*

Divergence time estimates indicate a crown age for *Sinocyclocheilus* of 15.08 Ma (95% HPD: 10.21–21.40 Ma, Figure S7). The LTT plot (Figure 4A) indicates a deviation from constant diversification rate in *Sinocyclocheilus*, which is also supported by the MCCR gamma statistics significantly rejecting constant diversification (γ = 2.1562; p < 0.05). This deviation is characterized by an acceleration in lineage accumulation since the group’s origin, with two notable peaks observed during the late Miocene, around 10.05 Ma and 8.56 Ma, followed by a slowdown beginning approximately at 8.10 Ma. Moreover, the LTT plot also identifies three surges in lineage accumulation during the Pleistocene, at 2.49 Ma, 0.53 Ma, and 0.09 Ma.

**FIGURE 4.**
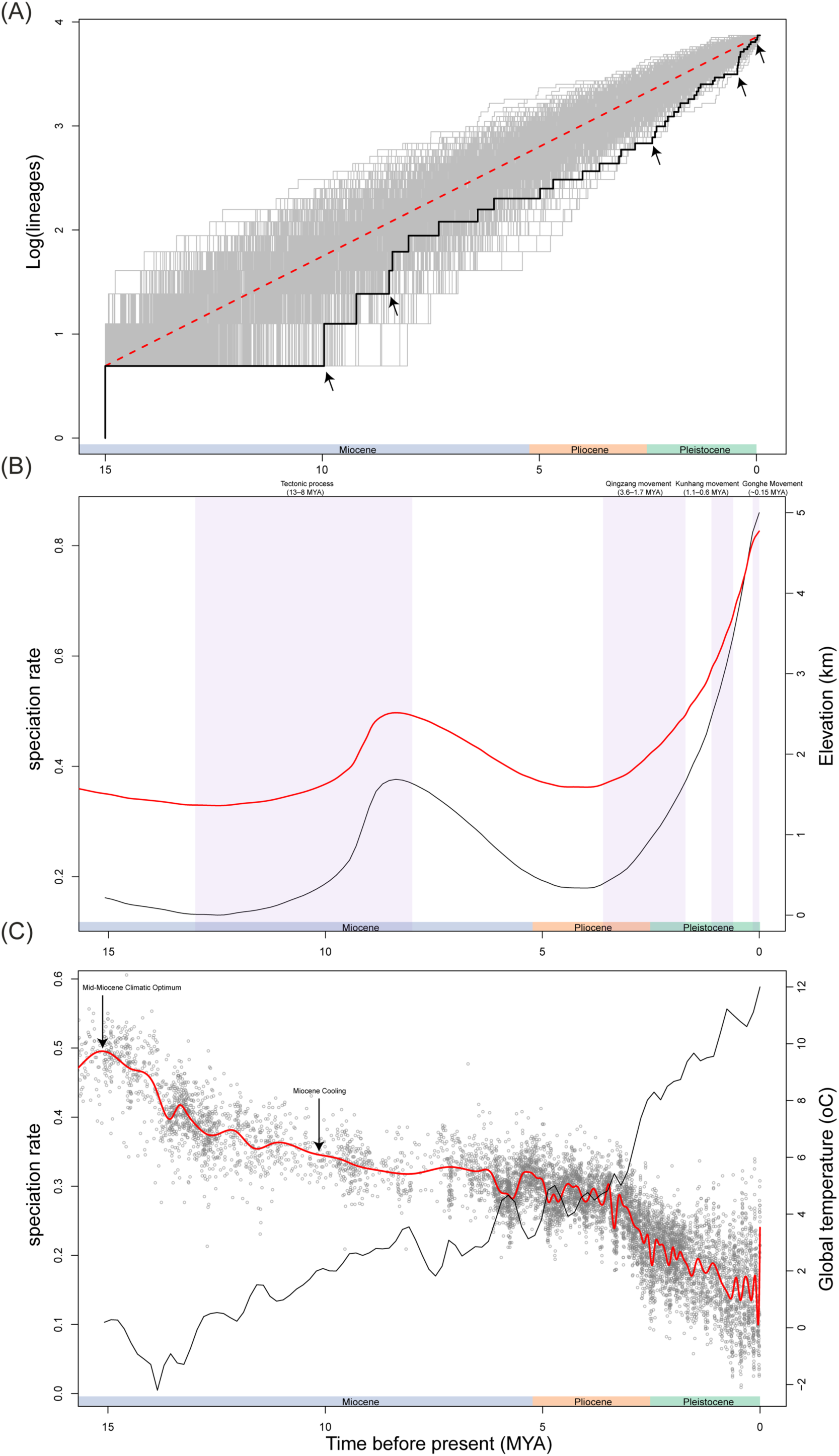
Diversification patterns of *Sinocyclocheilus*. (A) Observed LTT plot for *Sinocyclocheilus* based on the MCC tree (black line) compared to 1,000 simulated LTTs under a pure-birth process. The red dashed line shows the expectation from constant diversification through simulations, including the 95% confidence interval (gray area). Arrows indicate surges in lineage accumulation. (B) Speciation rates over time (black line) based on a model with linear dependence on paleo-elevation and zero extinction rates. The red line represents paleo-elevational estimates of the Tibetan Plateau. Periods of intense uplift (Qingzang, Kunhuang, and Gonghe movements) and tectonic processes are shaded in lilac. The decrease in elevation following the Gangdese and Himalaya Movements is attributed to prolonged erosion. (C) Speciation rates over time (black line) based on a model assuming linear dependence on paleo-temperatures and zero extinction rates. The red line represents global paleotemperature estimates.

The RPANDA analysis explored the relationship between the diversification of *Sinocyclocheilus* and various paleoenvironmental factors by fitting 18 models focused separately on paleoelevation of the Tibetan Plateau, global paleotemperature, and the intensification of East Asian monsoon (Table S7). The optimal model for paleoelevation demonstrated a positive linear relationship between speciation rates and the elevation of the Tibetan Plateau, with no associated extinction (AICω = 0.226, α = 0.213). This model suggests that speciation rates increased in parallel with the continued uplift of the Tibetan Plateau (Figure 4B). For paleotemperature, the best-fit model showed a negative linear relationship between speciation rates and historical temperatures, also with no extinction (AICω = 0.166, α = -0.08), indicating that speciation rates have risen as global temperatures decreased (Figure 4C). For monsoon effects, the most competitive model was the constant-rate speciation and extinction model (AICω = 0.179), while the second most competitive model (AICω = 0.109) revealed that increases in monsoon intensity are exponentially linked to higher speciation rates, along with a corresponding exponential increase in extinction rates (α = 2.021). This suggests that the speciation rate of *Sinocyclocheilus* may positively correlate with the East Asian monsoon.

### 3.4 State-dependent diversification and trait ancestral reconstruction

Using the HiSSE framework, we found that cave-adapted traits – degenerated eye, horn, hump, and reduced pigmentation – are significantly associated with diversification rates in *Sinocyclocheilus*. Specifically, the character-dependent HiSSE model ‘hisse.fit16’, characterized by reversible states, best explained the influence of horn and hump on diversification. Similarly, the character-dependent ‘hisse.fit12’ model, also featuring reversible states, was the best-fit model describing the impact of degenerated eye and reduced pigmentation on diversification (Table S8). The superior fit of these HiSSE models compared to the CID and BiSSE models suggests that diversification driven by these traits may be influenced by additional unmeasured factors, potentially including abiotic elements or a combination of intrinsic and extrinsic biotic factors. Trait-diversification plots indicate elevated diversification associated with each of the four traits evaluated (Figure 5). Specifically, a 6.7-fold increase in net diversification rates following transitions to a degenerated eye (Figure 5A), a 5.6-fold increase after transitions to horn presence (Figure 5B), a 6.7-fold increase after hump development (Figure 5C), and a 5.6-fold increase after transitions to reduced pigmentation (Figure 5D). Moreover, the best-fit of reversible models in the HiSSE model selection framework supports the reversibility of these four traits throughout the evolutionary history of *Sinocyclocheilus*.

**FIGURE 5.**
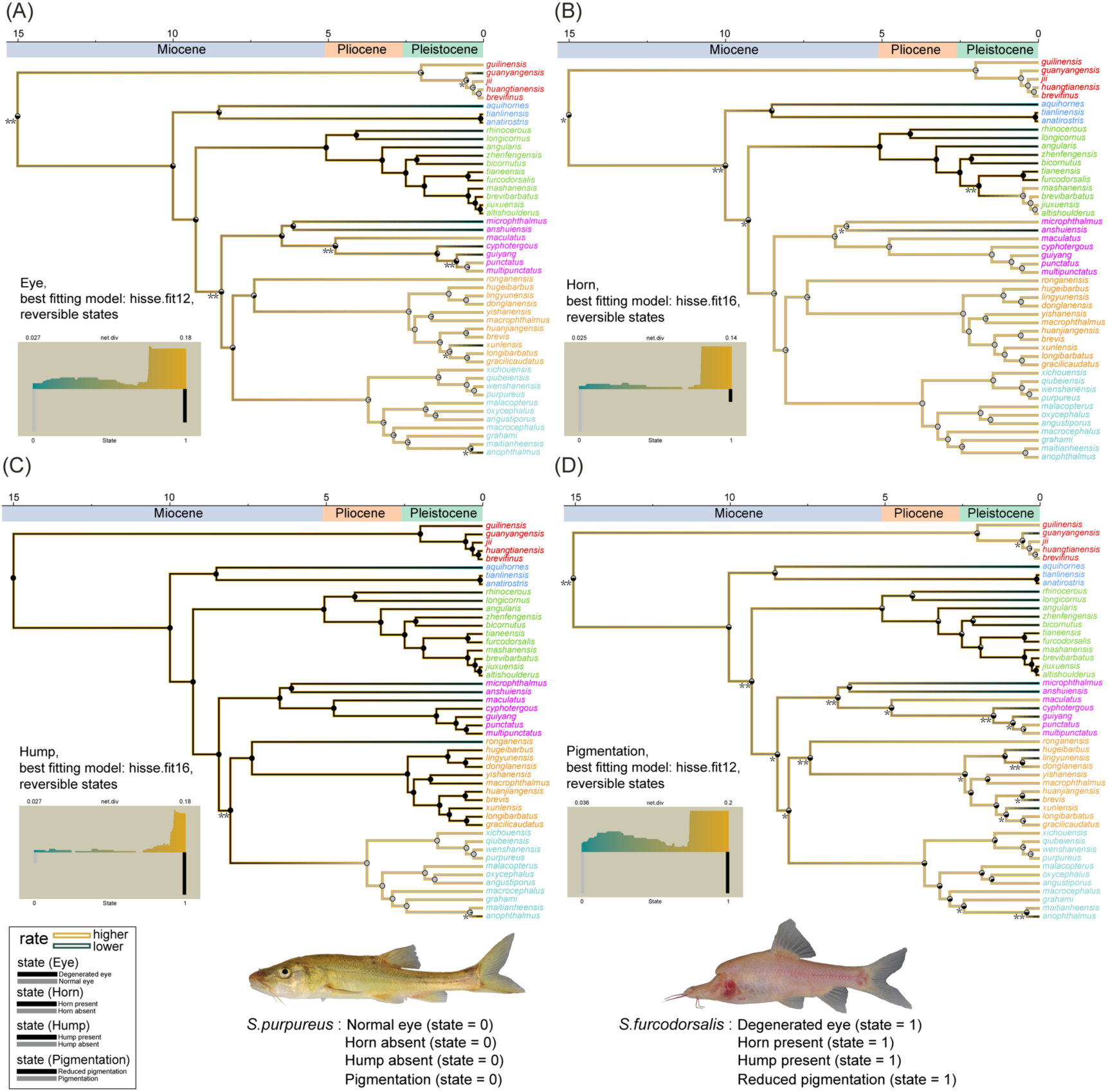
Phylogenetic reconstructions for four cave-adapted traits (degenerated eye, horn, hump and reduced pigmentation) in *Sinocyclocheilus* species. Net diversification rates and state reconstructions derived from the optimal HISSE trait-dependent model are mapped onto the chronogram. Ancestral state reconstructions with phytools are also mapped as pie-charts representing the probability of ancestral state onto the nodes of the chronogram. (A) Eye state reconstruction: transitions to degenerated eye marked with an asterisk (*), reversals to normal eye marked with a double asterisk (**). (B) Horn state reconstruction: transitions to horn presence marked with an asterisk (*), reversals to horn absence marked with a double asterisk (**). (C) Hump state reconstruction: transitions to hump presence marked with an asterisk (*), reversals to hump absence marked with a double asterisk (**). (D) Pigmentation state reconstruction: transitions to reduced pigmentation marked with an asterisk (*), reversals to pigmentation marked with a double asterisk (**). The phylogeny inner branches are color-coded to represent the reconstructed states, while the outer branches are shaded to indicate the net diversification rate, with the highest rates in gold and the lowest in dark cyan. Each panel’s lower corner features histograms displaying the distribution of net diversification rates for the observed states. These trait-dependent models imply an association between unobserved (hidden) states and the traits under study, which appear to influence the diversification process in *Sinocyclocheilus*.

The ancestral state reconstruction for degenerated eye, horn, hump, and reduced pigmentation favored the equal-rates model based on its superior Akaike Weight (Table S9). This model suggests the most probable ancestral states for the genus were degenerated eye (posterior probability = 0.508), absence of horn (posterior probability = 0.620), presence of hump (posterior probability = 0.989), and reduced pigmentation (posterior probability = 0.502) (Figure 5). Our ancestral reconstructions also indicate that both eye degeneration, and horn presence evolved independently three times, hump evolved only once, and reduced pigmentation evolved independently nine times (Figure 5). Conversely, the normal eye phenotype re-emerged at least four times, the horn phenotype was lost twice, hump lost once, and pigmentation reappeared seven times (Figure 5). These findings suggest that the evolution of these cave-adapted traits is a highly dynamic and reversible process.

### 3.5 Demographic history

The demographic histories of the 15 *Sinocyclocheilus* species, analyzed using Stairway Plot 2, (Figure 6; Figure S8) show a significant increase in effective population size (Ne) for each species during the late Pleistocene. Two main patterns emerged: the first pattern, evident in all species except *S. lingyunensis*, involved a rapid expansion followed by a period of stable effective population size (Ne) until after the Last Glacial Maximum (LGM), with multiple subsequent declines in Ne leading to the present (Figure 6). The second pattern, unique to *S. lingyunensis*, which exhibits partial cave-adapted traits such as a hump and reduced pigmentation, but retains normal eye and lacks a horn, experienced a population bottleneck around 150,000 years ago, followed by the most substantial population expansion observed among the species studied, with a stable Ne maintained until the present (Figure 6; Figure S8).

**FIGURE 6.**
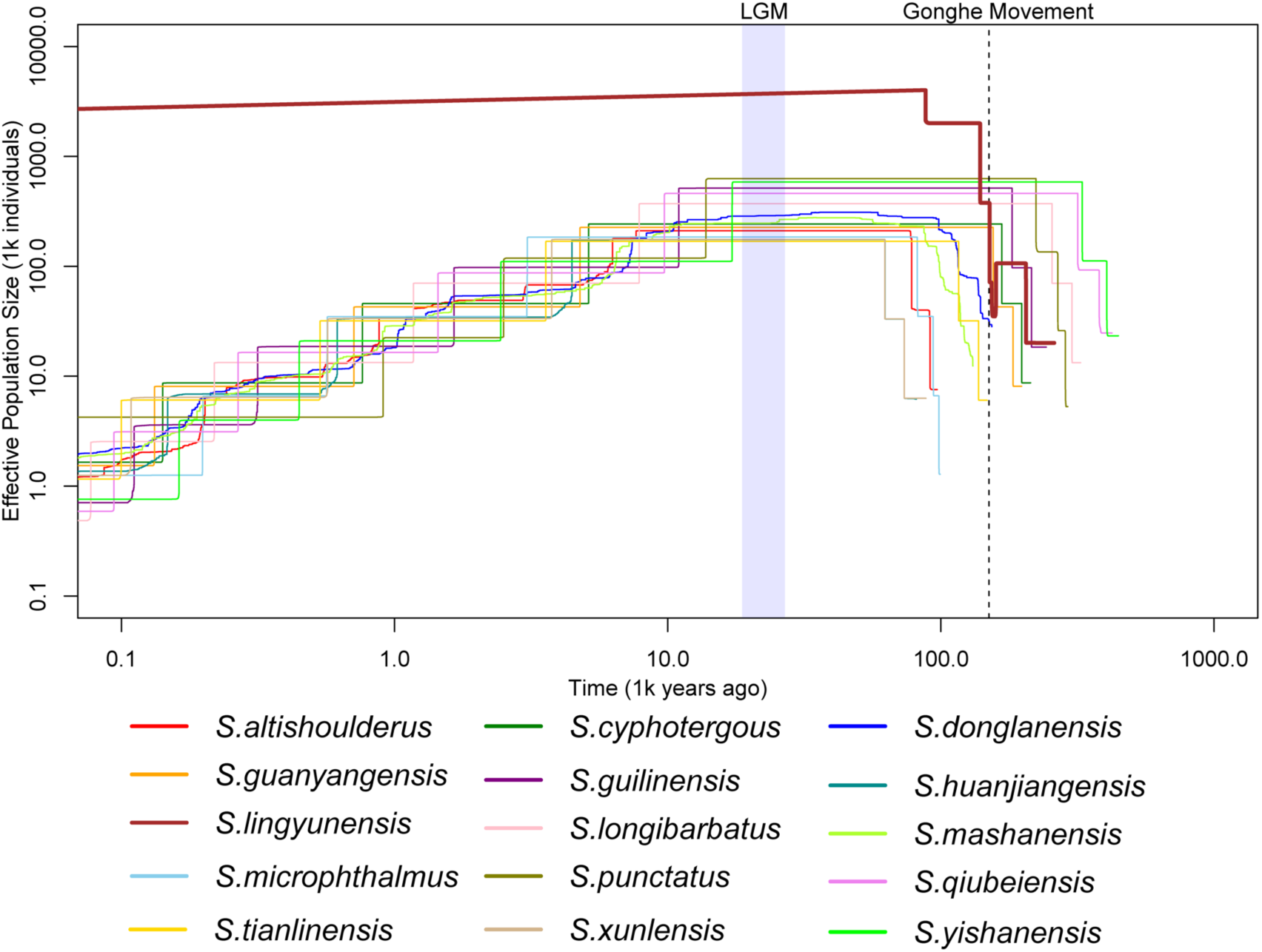
Demographic histories inferred by Stairway Plot2 for 15 *Sinocyclocheilus* species. The lilac vertical bar indicates the LGM period from 19 to 26.5 ka, while the vertical dashed line identifies the start of the most recent intense uplift of the Tibetan Plateau, known as the Gonghe movement, which occurred 150 ka ago.

## 4 DISCUSSION

*Sinocyclocheilus* is currently recognized as the largest genus of cavefish in the world. In this study, we developed an enhanced phylogenomic framework to investigate the biotic and abiotic factors driving diversification within this group, with a special focus on the roles of introgression, key innovations, geological events, and climate change. Our findings reveal widespread introgression within *Sinocyclocheilus*, which is closely linked to phylogenetic discordance and adaptive diversification. Through various analytical approaches, we identified the uplift of the Tibetan Plateau and global cooling as likely primary abiotic drivers of the rapid diversification in the genus.

Additionally, our results indicate that cave-associated phenotypes such as eye degeneration, horn development and reduced pigmentation are key innovations significantly contributing to diversification in this group.

Furthermore, the demographic history of several species appears to be shaped by the uplift of the Tibetan Plateau (particularly the Gonghe Movement), suggesting also that populations remained stable during the LGM, likely due to caves serving as glacial refugia.

### 4.1 Phylogenetic Discordance Reflects Introgression

*Sinocyclocheilus* cavefish, which has been the focus of recent research in evolutionary biology (T. Mao et al., 2022; T.-R. Mao et al., 2021; Wen et al., 2022; N. Yang et al., 2021), is thought to have undergone rapid diversification, with several cases of introgression reported (T. Mao et al., 2022). Unfortunately, disentangling phylogenetic relationships in cases of rapid radiations remains challenging due to the confounding effects of ILS and hybridization (Alexander et al., 2017; Astudillo-Clavijo et al., 2023; Degnan & Rosenberg, 2009; Maddison & Knowles, 2006; Roycroft et al., 2020). Our phylogenetic results reveal conflicts between gene trees and the species tree (Figure S3A, Figure S3B), as well as widespread discordance between the nDNA and mtDNA trees (Figure 1B). Our Dsuite, Treemix, and PhyloNetworks analyses indicate significant introgression between species pairs associated with these phylogenetic conflicts (Figure 2, Figure S5, Figure S6), suggesting that the discordance was likely driven primarily by introgression. The results from MSCquartets further support the conclusion that introgression largely contributes to these phylogenetic conflicts (Figure S4).

### 4.2 Pervasive Introgression and Natural Hybridization in Sinocyclocheilus

Introgression is prevalent and often facilitates adaptation and diversification in nature (Dowling & Secor, 1997; Edelman & Mallet, 2021). Numerous studies have identified cases of speciation with introgression, including examples in coral reef fish (Harrison et al., 2017), *Cyprinodon* pupfishes (Richards & Martin, 2017), cichlid fish (Salzburger, 2018), and *Triplophysa* fish (Qian et al., 2023). In this study, we employed three different methods to infer introgression among *Sinocyclocheilus* species. Each method revealed different patterns of gene flow, complementing one another, which could be attributed to the complexities of recurrent hybridization over varying time frames as well as the different sensitivities of these methods to gene flow intensity at different scales (Malinsky et al., 2018). Our findings demonstrate that ancient and recent introgression is widespread in this genus (Figure 2, Figure S5, Figure S6) and has occurred more frequently during the diversification of *Sinocyclocheilus* than suggested by earlier studies (T. Mao et al., 2022), indicating that gene flow may not necessarily impede speciation events. However, to the best of our knowledge, the isolation of caves or subterranean waters has likely limited genetic exchange between fish populations living in different caves (Ma et al., 2019). It is possible that in the cave-dense southern regions, where high temperatures, abundant precipitation, surface erosion, and fragile limestone formations prevail, frequent cave collapses or underground river diversions may have promoted secondary contacts between species. This could account for the extensive gene introgression observed. During the rapid diversification of *Sinocyclocheilus*, there may not have been enough time for the establishment of strong reproductive barriers, which might have been further weakened by repeated episodes of introgression. These frequent secondary contacts likely increased the potential for effective hybridization between *Sinocyclocheilus* species.

Research suggests that hybridization with ancestral lineages and genetic exchange between diverging lineages within a radiation could favor adaptive diversification (Marques et al., 2019b; Seehausen, 2004, 2013; Svardal et al., 2020). Our data reveal significant ancestral gene flow both between and within different *Sinocyclocheilus* clades (Figure 2C; Figure S5; Figure S6), suggesting that numerous hybridization events occurred between different *Sinocyclocheilus* ancestral lineages. Such ancient hybridization events could provide a reservoir for adaptive genetic variation (Marques et al., 2019b). Available ecological niche space allows selection to act upon genetic diversity arising from these ancient allele combinations, leading to the development of adapted and divergent phenotypes (Seehausen, 2004, 2013). Consequently, these hybridization events may have fueled the radiation and diversification of *Sinocyclocheilus*.

The Syngameon hypothesis (Gillespie et al., 2020; Seehausen, 2004) posits that hybridization acts as a creative force, enabling hybrid lineages to explore new trait spaces—a critical factor in adaptive radiations. Greater divergence between species prior to hybridization increases the potential for hybridization to introduce novel and advantageous recombinant genotypes and structural variants, which can lead to phenotypic innovation (Seehausen, 2004; Stelkens & Seehausen, 2009). Our findings provide evidence of introgression between different ecomorphs, such as between the surface species *S. ronganensis* and the cave species *S. aquihornes* (Figure S6A), between the surface species *S. punctatus* and the cave species *S. cyphotergous* (Figure 2), and between the surface species *S. longibarbatus* and the cave species *S. xunlensis* (Figure 2). These observations support the Syngameon hypothesis of adaptive radiation, which argues that hybridization between non-sister lineages within an adaptive radiation facilitates further ecological divergence and speciation (Gillespie et al., 2020; Seehausen, 2004).

The impact of hybridization in facilitating adaptive radiation hinges on whether it leads to hybrid lineages evolving more rapidly or accessing novel trait spaces, either through unique combinations of parental traits or by developing trait values beyond the range of their parental lineages (Parsons et al., 2011; Seehausen, 2004). In this study, ongoing hybridization is observed between the sympatrically distributed cave species *S. cyphotergous* and the surface species *S. punctatus*, supported by the identification of two hybrid individuals (Liu129 and Liu354), which display intermediate phenotypes and significant genomic admixture (Figure 3). Notably, the individual Liu354 displays cave-adapted traits, such as degenerated eye, reduced pigmentation, and a pronounced hump, distinguishing it from the normal phenotype of the surface species *S. punctatus* (Figure 3B). Our results suggest that hybridization between cave and surface species may contribute to localized variation in traits associated with cave adaptation, including eye development, pigmentation, and hump formation, which may drive ecological diversification in the adaptive radiations of *Sinocyclocheilus*.

### 4.3 Geographic and Climatic Factors Driving the Radiation of Sinocyclocheilus

The remarkable species diversity within *Sinocyclocheilus* has been linked to major geological events, particularly the intense uplift of the Tibetan Plateau and the distribution of karst underground rivers (S. Chen et al., 2009; Xiao et al., 2005). However, no prior studies have explicitly modeled the diversification dynamics of *Sinocyclocheilus* in relation to the uplift of the Tibetan Plateau over time. Our LTT plot (Figure 4A) revealed five significant surges in lineage accumulation, which closely align with tectonic events in the Tibetan Plateau. Employing a paleoelevation-dependent diversification model, we found a strong correlation between the speciation rate in *Sinocyclocheilus* and the uplift of the Tibetan Plateau (Figure 4B; Table S7). The frequent capture and isolation of subterranean river systems, combined with karst development, are considered key drivers of speciation in this genus (Zhao & Zhang, 2009). The rapid uplift of the Tibetan Plateau has resulted in extensive and continuous fragmentation of both surface and subsurface fluvial habitats, providing ecological opportunities and geographic isolation, which likely contributed to the diversification of *Sinocyclocheilus*.

Adverse environmental conditions, such as habitat loss, increased cold, or aridity, can transform caves into refuges for troglophilic species inhabiting both hypogean and epigean environments (Bryson et al., 2014; Camp et al., 2014; Culver, 1982; Mammola et al., 2019). Using a paleotemperature-dependent diversification model, we found that the speciation rates in *Sinocyclocheilus* were positively associated with global cooling (Figure 4C; Table S7). During cooler periods, many *Sinocyclocheilus* species likely could not tolerate the harsh surface conditions and repeatedly sought refuge in stable karst cave waters. These caves, with stable, isothermal water temperatures (Howarth, 1980; Tabin et al., 2018), likely provided ideal refuges, promoting population isolation and, consequently, promoting diversification. Furthermore, climatic fluctuations during the Pleistocene, alternating between cold and warm periods, may have led to repeated ingress and egress of *Sinocyclocheilus* from caves, facilitating secondary contact. Our results also suggest a positive relationship between diversification rate and monsoon intensity (Table S7). The intensification of the East Asian monsoon since the Miocene, characterized by increased precipitation, likely enhanced karst development (Kong et al., 2017). Active karstification, which led to the formation of karst caves, likely provided suitable habitats and favorable geographical conditions for the early diversification of *Sinocyclocheilus*.

### 4.4 Four Cave-adapted Traits are Associated with Rapid Diversification Rates

Caves present harsh environments characterized by perpetual darkness and limited food availability (Borowsky, 2018; White & Culver, 2012). *Sinocyclocheilus* cavefish have undergone both constructive and regressive morphological adaptations to thrive under these challenging conditions (Ma et al., 2019). Our HiSSE analyses suggest that the development of constructive traits, such as the horn and hump, in combination with additional, unmeasured factors, may have contributed to increased net diversification rates in *Sinocyclocheilus* (Table S8). To cope with the scarcity of environmental food resources, troglobitic cavefish must optimize energy storage (Fišer, 2019; Riddle et al., 2018; Xiong et al., 2022). The hump, composed primarily of adipose tissue lacking bone structures, serves as an energy reservoir (Lunghi et al., 2019). The horn, comprising the frontal and parietal bones, may store fat to nourish the brain (Ma et al., 2019). Additionally, the horn likely functions as a complex sensory organ, partially compensating for the sensory roles of barbels and the lateral line (Y.-R. Chen et al., 1994; Soares et al., 2019), while also providing a protective structure to shield the brain from potential damage caused by collisions with cave walls (W. Li & Tao, 2002; Soares et al., 2019).

In addition to increasing energy storage, cave-adapted species also strive to minimize energy expenditure (Hervant, 2012; D. Moran et al., 2014, 2015; J.-H. Zhang et al., 2023). Our HiSSE analyses revealed that regressive traits, such as eye degeneration and reduced pigmentation, along with additional unmeasured factors, may have also contributed to the elevated net diversification rates observed in *Sinocyclocheilus* (Table S8). The degeneration of eyes, which have high energy demands, allows cavefish to conserve up to 15% of their energy (Krishnan & Rohner, 2017). In contract, pigmentation in surface-dwelling organisms, serves various functions such as protection from sunlight, camouflage, mimicry, and species or sex recognition (Protas & Patel, 2008)—all of which are irrelevant in the perpetual darkness of caves.

Cavefish adaptations generally follow a common theme of eliminating nonessential morphological traits to minimize energy expenditure while promoting the development of organs that increase energy reserve capacity. These adaptations include eye degeneration, reduced pigmentation, and the development of structures such as horn and hump. Together, these changes confer a selective advantage in cave environments with irregular food supplies. Additionally, HISSE models suggest that other traits might potentially underpin diversification regimes within *Sinocyclocheilus* (Table S8). These include various constructive and regressive morphological changes such as protruding jaw, increased numbers of taste buds, overdeveloped barbels, loss of diurnal rhythms, potential hearing loss, and scale reduction. These features warrant further investigation to determine their roles in contributing to cave adaptation and the diversification of *Sinocyclocheilus* cavefish.

### 4.5 Transitions of Cave-adapted Traits in *Sinocyclocheilus*

Previous ancestral reconstructions using mtDNA phylogeny suggested that the ancestral *Sinocyclocheilus* had degenerated eyes and lacked a horn (T.-R. Mao et al., 2021). Our reconstruction, based on RAD-based phylogeny, aligns with these findings, confirming that the ancestral *Sinocyclocheilus* indeed possessed degenerated eyes and lacked a horn (Figure 5). Furthermore, we discovered that the ancestral state included the presence of hump and reduced pigmentation, reinforcing the notion that *Sinocyclocheilus* exhibited pre-adaptations for cave dwelling. Our ancestral state reconstruction analyses further indicated that nearly all transitions to cave-adapted traits occurred independently during the late Miocene and Pleistocene (Figure 5), coinciding with the uplift of the Tibetan Plateau and global cooling events. Environmental fluctuations in the ecogeographical landscape associated with the Tibetan Plateau uplift, may have driven the cyclical loss and re-gain of adaptive morphological traits throughout the history of *Sinocyclocheilus* cavefish in the karst region.

### 4.6 Demographic History of 15 *Sinocyclocheilus* Species

Previous research identified a population expansion event in *S. rhinocerous* during the interglacial period (0.5–0.15 Ma), attributed to the extensive subterranean water systems in its native Luoping Basin (J. Yang et al., 2016). In our demographic reconstruction, we observed similar population expansion events during this period in eight *Sinocyclocheilus* species (Figure 6). This suggests that extensive subterranean water systems may have been prevalent in their habitats as well.

Environmental fluctuations and substantial hydrographic changes may have facilitated the contiguous range expansion of freshwater fish populations, providing opportunities for increased population sizes in some species (Deng et al., 2021; Su et al., 2015; G. Wang et al., 2022; L. Yang et al., 2009). The reconstruction of the demographic history for *Sinocyclocheilus* species suggests that following the Gonghe Movement, eight species underwent population expansions (Figure 6), possibly due to the dramatic tectonic and climatic events that significantly impacted the current drainage systems (Hou et al., 2012; D. Liu et al., 2015; Zhou et al., 2018). These events likely led to the expansion of the underground river system and enhanced connectivity, facilitating these changes.

Glacial refugia in southwestern China, including plateaus, basins, mountain peaks, and deep gorges, protected many organisms from the climatic fluctuations of the LGM (Fan et al., 2012; Fu & Wen, 2023; Z. Li et al., 2012; Song et al., 2018; Yu et al., 2013; Zheng et al., 2020). Our findings indicate that the effective population size remained stable across all 15 *Sinocyclocheilus* species during the LGM (Figure 6). Unlike other glacial refugia, caves likely served as refuges for *Sinocyclocheilus*, providing isolated yet climate-stable habitats that mitigated the adverse effects of the LGM. Following the LGM, 14 out of 15 *Sinocyclocheilus* species experienced a population decline (Figure 6), which may be attributed to the multiple episodes of aridification experienced in southwest China during this period (Y. Li et al., 2009; X. Wang et al., 2023; W. Zhang et al., 2018).

Introgression can facilitate adaptive changes in response to environmental shifts (Brauer et al., 2023; De-Kayne et al., 2022; Ge et al., 2023). *S. lingyunensis* is the only species that maintained a stable effective population size after the LGM (Figure 6). This stability may be attributed to significant genetic introgression between *S. lingyunensis* and other *Sinocyclocheilus* species (Figure 2), which may help mitigate threats such as aridification.

Additionally, deleterious alleles are often eliminated from population through random genetic drift, particularly in populations with a small effective population size (Ne). Gene flow plays a crucial role in masking these deleterious alleles, thereby supporting the sustained growth rate of populations (Hoffmann et al., 2021; Khan et al., 2021; Qian et al., 2023; Whiteley et al., 2015).

## 5 CONCLUSIONS

Our study shows that introgression, rather than incomplete lineage sorting, significantly contributes to the phylogenetic discordance observed in *Sinocyclocheilus*. The rapid diversification of these species appears to be bolstered by pervasive introgression, as evidenced by the observed hybridization between sympatric species, with hybrid individuals exhibiting intermediate phenotypes of their parental species. This underscores the significant role of hybridization in the evolutionary history of *Sinocyclocheilus*. Additionally, the uplift of the Tibetan Plateau and global cooling likely played a major role in driving rapid species diversification, while the intensification of the East Asian monsoon may have only indirectly influenced this diversification. Three key innovations in *Sinocyclocheilus* cavefish—eye degeneration, horn, and reduced pigmentation—facilitated the adaptation to dark cave environments and also promoted the diversification of *Sinocyclocheilus*. After the Gonghe Movement, many Sinocyclocheilus species experienced population expansions, followed by relative stability during the Last Glacial Maximum (LGM), with caves serving as glacial refugia.

Our study demonstrates the contributions of both biotic (hybridization and key innovations) and abiotic factors (ecological opportunities provided by climate and/or geological changes) in the diversification of *Sinocyclocheilus* cavefish. We highlight the importance of integrating biotic and abiotic drivers, rather than focusing on a single factor, to fully explain the macroevolutionary patterns of this radiating lineage.

## AUTHOR CONTRIBUTIONS

TRM, MM, MMV, MRP, YWL and JY conceptualized the research and designed the methodology; YWL, TRM, MM, JY and SPZ conducted fieldwork and curated the data; TRM, MMV, MRP, GE and MM carried out formal analysis; TRM, MM, MMV and MRP wrote the original draft; TRM, MM, MMV, MRP, SPZ, GE and YWL made figures; MM, MMV, and MRP supervised; All authors reviewed and edited the draft. All authors read and approved the final manuscript.

## ACKNOWLEDGEMENTS

We thank the EED lab members for fieldwork; Cheng-Hai Fu for field work; Rohan Pethiyagoda (Ichthyology Section, Australian Museum) for commenting on the manuscript.

## ETHICS STATEMENT

The animal study was reviewed and approved by the Institutional Animal Care and Use Committee of Guangxi University (GXU), Nanning-China (#GXU2019-071). Field sampling was approved by the Guangxi Province Government; fish were sampled using trap nets, and tissues from the fish were obtained using fin clips without destructive sampling.

